# N6-methyladenosine (m^6^A) RNA modification restrains antifungal immunity and is a therapeutic target in oral candidiasis

**DOI:** 10.1101/2025.09.07.674635

**Authors:** Nicole O. Ponde, Melissa E. Cook, Isha Mehta, Ipsita Dey, Kenta Yamamoto, Tiffany C. Taylor, Bianca M. Coleman, Shachi P. Vyas, Rami Bechara, Jishnu Das, Sarah L. Gaffen

## Abstract

Fungal infections represent a major global threat, yet to date there is a paucity of effective antifungal drugs and no licensed vaccines to any pathogenic fungi. The commensal pathobiont *Candida albicans* can cause severe oropharyngeal and mucocutaneous infections in immunocompromised individuals. The initial point of interaction of *C. albicans* in the oral mucosa is with superficial oral epithelial cells (OECs). Upon fungal encounter, OECs upregulate a program of anti-fungal defense genes, a process that has been extensively studied at the level of proximal signaling pathways, transcriptional induction, and epigenetic control. However, inflammatory genes are also subject to extensive regulation at the mRNA level, yet little is known about mechanisms that control *C. albicans*-induced genes post-transcriptionally. The importance of ‘epitranscriptomic’ gene regulation by mRNA methylation is becoming increasingly appreciated, yet its contribution to host responses during fungal infection remains poorly understood. N6-methyladenosine (m^6^A) is the most abundant RNA modification and influences numerous transcript fates including mRNA stabilization. Here, we demonstrate that *C. albicans*-induced genes in human and mouse OECs are widely subject to m^6^A modification following infection. Silencing of a specific subclass of m^6^A binding proteins, the YTHDF1/2/3 ‘readers’, reprogrammed the OEC response to *C. albicans* by up- and down-regulating antifungal transcripts. The net effect of blocking the m^6^A pathway using a first-in-class METTL3 inhibitor was to amplify mRNA expression of antimicrobial peptides that reduce fungal burden. These studies establish a role for the m^6^A pathway as a modulator of immunity to mucosal candidiasis and suggest a new therapeutic avenue to treating this condition.

## Introduction

Fungal infections represent a significant yet under-appreciated contributor of morbidity and mortality in humans. This is underscored by the designation of *Candida albicans* (*C. albicans*) as a WHO priority pathogen together with a lack of licensed vaccines and a paucity of approved antifungal therapies. *Candida* species are a leading cause of human fungal infections, encompassing both superficial and invasive diseases. *C. albicans* is the most common causative agent, implicated in a wide array of clinical presentations, including oral, vaginal, and systemic infections ^1^.

Oropharyngeal candidiasis (OPC, oral thrush) varies from mild, self-resolving infections to severe, painful conditions that can be associated with nutritional deficits and esophageal cancer. In the mouth, the first cells to encounter *C. albicans* are epithelial/keratinocyte cell types, and this population is a central coordinator of early responses to these fungi. Superficial oral epithelial cells (OECs) line mucosal surfaces including tongue, palate and buccal mucosae, sites that are characterized by continuous microbial exposure and rapid epithelial cell turnover and renewal ^2^. Interactions between *C. albicans* and OECs initiate a complex signaling cascade that drives effective host defense ^3–6^. Ultimately, integration of these sensing pathways culminates in transcriptional induction of immune-related genes, which include proinflammatory cytokines, chemokines, antimicrobial peptides (AMPs), and other effectors that control infection yet restrain immunopathology ^5,7–10^.

While transcriptional regulation is a critical facet of the host response to *C. albicans* infection ^5,6,11^, post-transcriptional responses to *Candida* signaling are surprisingly poorly understood. Inflammatory gene transcripts are highly subject to control not only through *de novo* transcription but also through multiple post-transcriptional mechanisms, including mRNA stability, control of translation, RNA nuclear export, splicing, etc. ^12–19^.

Among the many mechanisms governing post-transcriptional gene regulation, epitranscriptomic modifications have emerged as determinants of host defenses and cellular function. By influencing RNA processing, translation, and decay, modifications of RNA enable rapid or nuanced changes in gene expression that are independent of the underlying genetic sequence. N6-methyladenosine (m⁶A) is the most abundant RNA modification in eukaryotic cells, controlling diverse eukaryotic processes including metabolism, inflammation and host defense ^20–23^. m^6^A is installed predominantly within a conserved DRA*CH motif (A* is the methylatable adenosine, D=A, G or U; R=A and G; H=A, C or U). The m⁶A modification occurs at an average of ∼3 sites per mRNA within the consensus DRACH motif, distributed across coding sequences and UTRs of several thousand transcripts ^24–28^. Such modifications are typically enriched in 3’ untranslated regions (UTRs), long internal exons, or proximal to stop codons. Deposition of m^6^A is conferred by methyltransferases (termed ‘writers’; METTL3, METTL14) and can be reversed by demethylases (’erasers’; FTO, ALKBH5), allowing for dynamic control of this process ^20^.

The functional consequences of m^6^A modification are dictated by ‘reader’ RNA binding proteins (RBPs), which recognize m^6^A directly or bind to sequence structures influenced by its presence ^20,23,28–31^. Among the best characterized m⁶A reader subclasses are the YT521-B homology (YTH) domain-containing proteins and the insulin-like growth factor 2 (IGF2BP) family ^32,33^. The YTHDF family (YTHDF1–3) readers have emerged as major regulators of RNA fate, strongly influencing the magnitude and duration of gene expression programs ^34–36^. Although understanding the functions of individual YTHDF family members remains an active area of inquiry ^36,37^, YTHDFs have generally been found to destabilize target mRNA transcripts ^38,39^, positioning them as post-transcriptional regulators that constrain inflammation. However, the outcome of m⁶A deposition is not universally suppressive. The IGF2BP family of m^6^A readers (also known as IMPs) are best characterized to stabilize methylated transcripts and/or to promote RNA translation ^32,40^.

There is growing interest in how m⁶A-dependent RNA regulation shapes immune responses during infection. Recent evidence indicates that RNA methylation pathways within *C. albicans* contribute to its pathogenicity, and pharmacological inhibition of fungal methyltransferase activity impaired virulence ^41^. Although there are demonstrated roles for m^6^A modification in viral and bacterial infections ^16,42–46^, the consequences of m^6^A modification in antifungal host defense are almost entirely underexplored. Here, we demonstrate that many mRNAs induced by *C. albicans* infection in the oral epithelium are highly methylated, and that YTHDF m^6^A readers influenced *Candida*-induced transcriptional programs. *In vivo*, blockade of the m^6^A pathway using a first-in-class pharmacological METTL3 inhibitor improved outcomes in murine oropharyngeal candidiasis, which was linked to enhanced antimicrobial protein expression and accelerated fungal clearance. Thus, on balance, the m^6^A pathway restrains inflammatory antifungal immunity and thereby promotes fungal pathogenicity. Together, these data highlight a role for epitranscriptomic regulation in host-pathogen interactions during *C. albicans* infection and suggest a pathway for drug targeting.

## Results

### YTHDF m^6^A readers regulate genes bidirectionally during *C. albicans* infection

To investigate whether the m^6^A pathway in any way influences host responses to *C. albicans* infection, we performed a targeted siRNA screen of m^6^A pathway writers, erasers and readers in TR146 cells, a human OEC line widely used to interrogate epithelial-specific oral defenses against *C. albicans* ^8,10,47–50^. Cells were transfected with siRNAs targeting the core m^6^A writer METTL3, the FTO and ALKBH5 demethylase erasers, and the 3 major classes of m^6^A readers (YTHDF, YTHDC and IGF2BP2 family members). Forty-eight hours after siRNA transfection, cells were cultured with *C. albicans* yeast (strain SC5314) for 4 h (**Fig. 1a**), a time frame that allowed *C. albicans* to transition to its virulent hyphal form, and expression of representative *C. albicans*-induced inflammatory genes (*CCL20, IL36*) were assessed ^10,50^. There were no detectable differences following knockdown of METTL3 or the m^6^A erasers (FTO, ALKBH5). There was also no impact of silencing the YTHDC and IGF2BP m^6^A reader families on these genes (**Fig. 1b**). However, silencing of the YTHDF family (YTHDF1, 2, 3) led to upregulation of *CCL20* and *IL36* (**Fig. 1 c)**. This result is in line with the well-characterized property of YTHDF readers to suppress target mRNA expression through RNA destabilization ^16,51–54^. Individual knockdown of YTHDF1/2/3 similarly enhanced target gene expression, suggesting redundancy among these factors (**Extended Data Fig. 1**). Although it was somewhat paradoxical that silencing of central m^6^A pathway effectors (METTL3 or m^6^A erasers) did not impact these target genes while knockdown of a specific reader family (YTHDFs) did, it is likely that the combined impact of other cellular effects mediated by m^6^A modifications served to offset the specific effects of YTHDFs. Thus, YTHDF m^6^A readers function as negative regulators of a subset of genes associated with epithelial antifungal defense.

**Figure 1.**
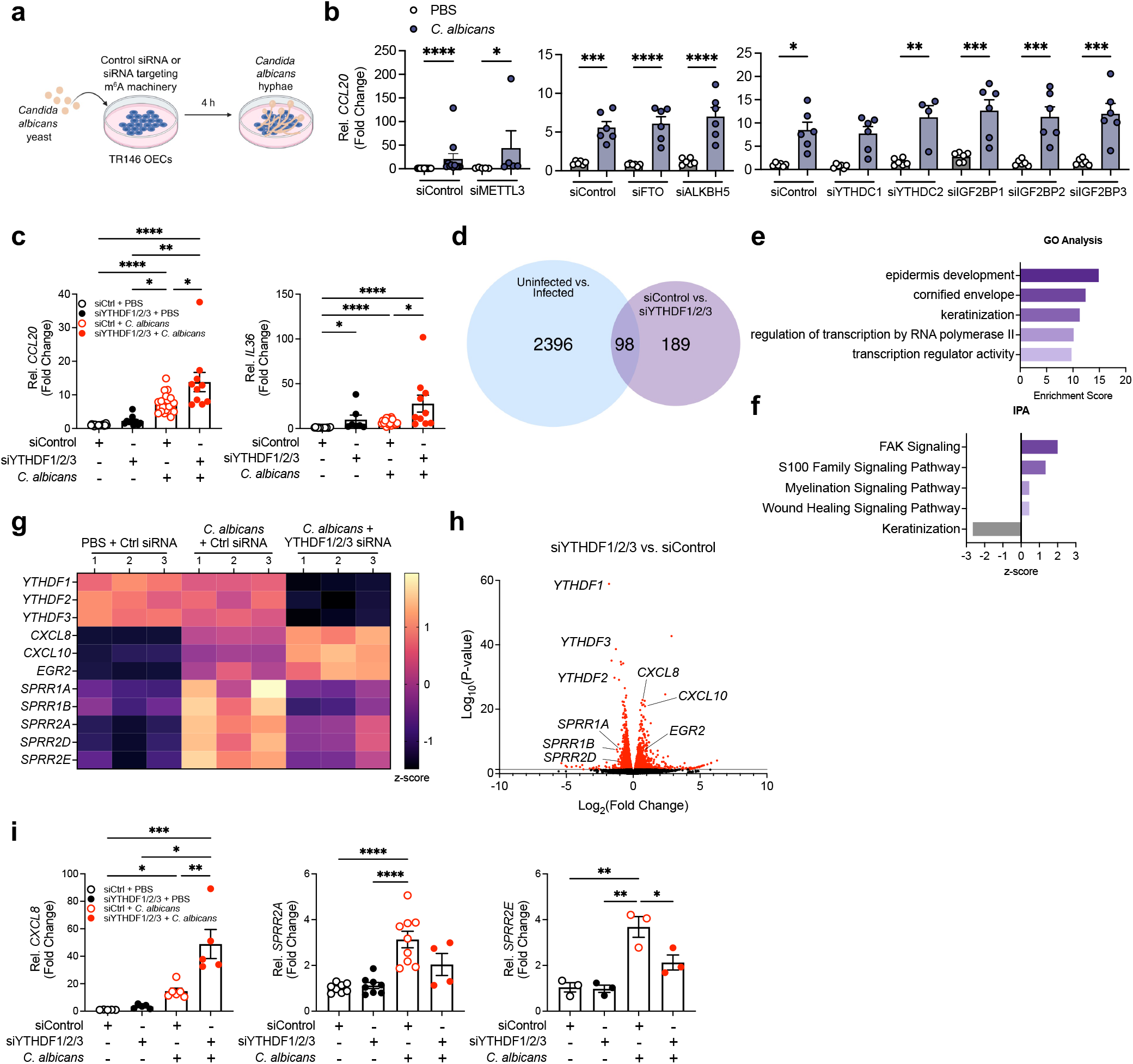
YTHDF readers bidirectionally regulate *C. albicans*-induced inflammatory responses in OECs. (**a**) Experimental overview. (**b-c**) TR146 cells were transfected with the indicated siRNAs and stimulated with *C. albicans* yeast (SC5314) 48 h post-transfection. *CCL20* was measured by qPCR, normalized to *GAPDH* and expressed as fold-change relative to siCtrl +PBS. Date show mean ± SEM (n=6). Significance was assessed by Kruskal-Wallis with Dunn’s multiple comparisons test. (**d**) TR146 cells were transfected with the indicated siRNAs, infected with *C. albicans* yeast, and total RNA was subjected to Illumina RNASeq (n=3 per group). DEGs that are suppressed or enhanced upon YTHDF1/2/3 knockdown and affected by *C. albicans* infection. (**e**) Gene ontology (GO) analysis (**f**) Ingenuity Pathway Analysis (IPA) of differentially regulated pathways. (**g, h**) *C. albicans*-induced genes differentially expressed upon YTHDF1/2/3 knockdown (n=3). (**i**) Validation of *CXCL8, SPRR2A* and *SPRR2E* following YTHDF1/2/3 knockdown (n= 3-8). Significance was assessed by 1-way ANOVA with Bonferroni’s multiple comparisons, Kruskal-Wallis with Dunn’s multiple comparisons or Mann-Whitney test.

To define the landscape of YTHDF-mediated gene regulation across the transcriptome, we performed RNA-seq in the same conditions as the initial screen (YTHDF1/2/3 silencing in TR146 cells followed by *C. albicans* exposure for 4 h). In total, 98 differentially expressed genes (DEGs) were identified as potential targets of YTHDFs, defined as intersecting set of genes affected by knockdown (siCtrl vs. siYTHDF1/2/3) and genes that are induced by *C. albicans* (uninfected vs. infected) (**Fig. 1d**). Gene ontology (GO) analysis and Ingenuity Pathway Analysis (IPA) indicated that pathways associated with keratinization, wound healing/FAK signaling, transcriptional regulation, and inflammation (S100 and myelination signaling) were strongly impacted by YTHDF silencing (**Fig. 1e-f**). Several canonical *C. albicans*-induced proinflammatory transcripts were upregulated, implying a suppressive effect of YTHDFs. These included chemokines *CXCL8* and *CXCL10* and the transcription factor (TF) *EGR2* ^9^ (**Fig. 1g-h**). However, some genes were suppressed following knockdown, notably members of the small proline-rich (SPRR) antimicrobial peptide/wound healing family that were recently shown to have direct anti-*Candida* activity (**Fig. 1g-h**) ^55,56^. Confirmation of selected genes corroborated the RNASeq results (e.g., *CXCL8* was elevated while *SPRR2A* and *SPRR2E* were reduced) (**Fig. 1i**). Together, these findings reveal that YTHDF m^6^A readers function as suppressors of certain inflammatory genes while promoting expression of others.

### *C. albicans*-induced YTHDF-regulated transcripts are subject to m⁶A modification

As molecular interpreters of m⁶A-marks, readers such as the YTHDFs bind preferentially to RNA that is methylated ^37,54^. Putative m⁶A consensus DRA*CH motifs were identified using the SRAMP prediction platform ^57,58^, which revealed methylation sites of varying confidence across most candidate transcripts (**Table 1**). To determine whether the genes in *C. albicans*-infected TR146 cells contained m⁶A modified nucleotides, lysates containing total RNA were immunoprecipitated with anti-m^6^A antibodies and subjected to qPCR (m⁶A RNA immunoprecipitation (MeRIP). As expected based on SRAMP predictions, *CXCL8* and *SPRR2A* were enriched within the MeRIP fraction following *C. albicans* stimulation as compared to sham (PBS-treated) samples and IgG pulldown controls (**Fig. 2a**).

**Figure 2.**
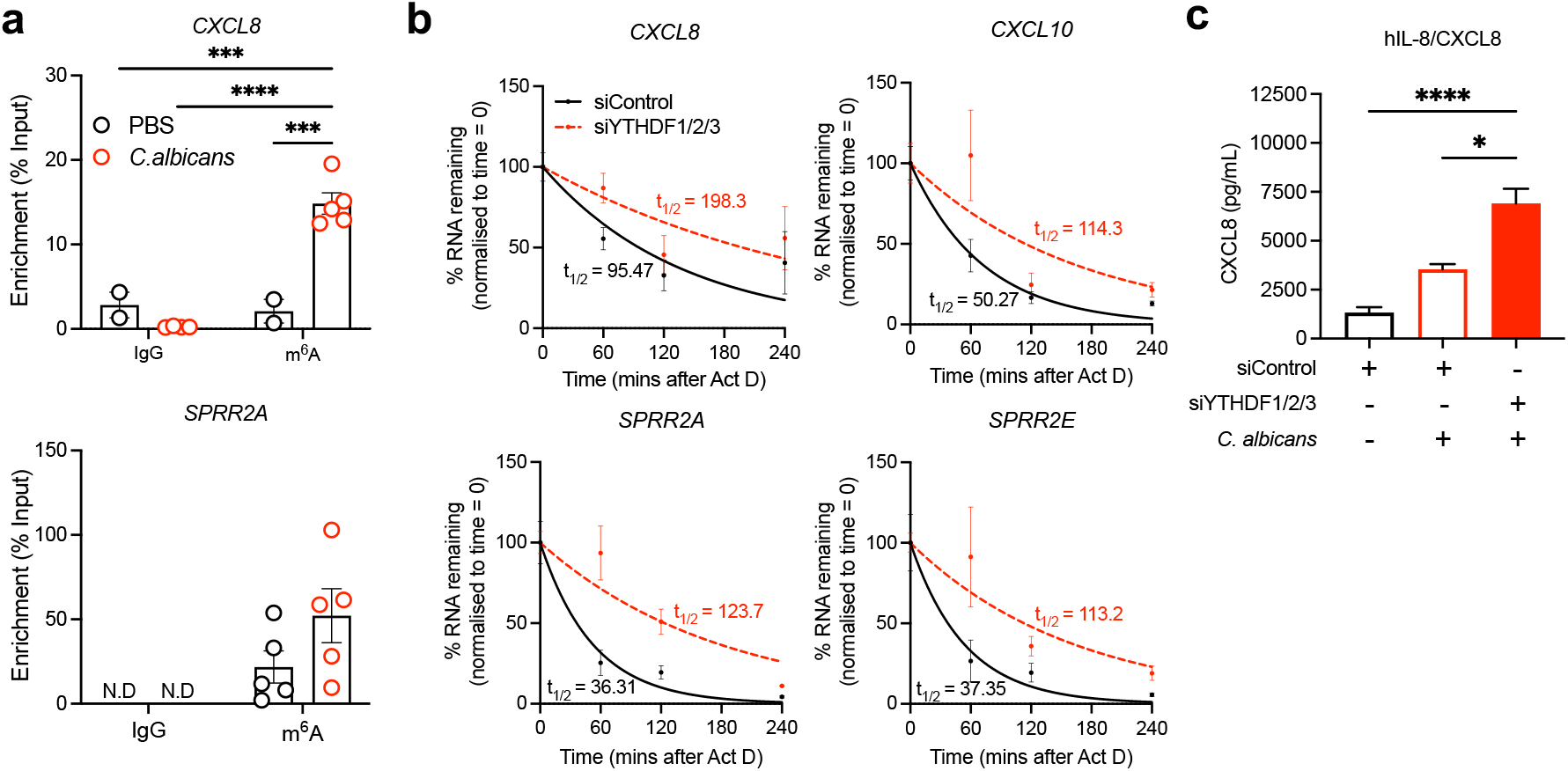
YTHDF readers regulate infection-induced transcripts. (**a**) TR146 cells were treated with PBS (black) or stimulated with *C. albicans* yeast (red) for 4 h. RNA was subjected to MeRIP and *CXCL8* and *SPRR2A* were quantified by qPCR, normalized to *GAPDH* (n= 3-5). Significance was assessed by 2-way ANOVA with Bonferroni’s multiple comparisons test. (**b**) TR146 cells were transfected with the indicated siRNAs and stimulated with *C. albicans* for 4 h. Cells were treated with actinomycin D and mRNA quantified by qPCR at the indicated times. Graphs depict the percentage of RNA remaining relative to the 0-min time point, with one phase decay curves assessed by non-linear regression (n=3). (**c**) TR146 cells were subjected to YTHDF1/2/3 silencing and infected with *C. albicans* for 8 h. CXCL8 in supernatants was assessed by ELISA (n=2). Significance was assessed by Kruskal-Wallis with Dunn’s multiple comparisons test.

**Table 1.** Predicted sites of m^6^A modification in the indicated transcripts, identified using the SRAMP database.

| Gene Name | SRAMP Score (Confidence) |  |  |  |
| --- | --- | --- | --- | --- |
|  | Very High | High | Moderate | Low |
| *CXCL8* | 1 | 2 | 3 | 1 |
| *CXCL10* | 0 | 0 | 3 | 1 |
| *CCL20* | 1 | 1 | 2 | 3 |
| *SPRR1A* | 0 | 1 | 1 | 1 |
| *SPRR2A* | 0 | 0 | 1 | 0 |
| *SPRR2D* | 0 | 0 | 0 | 2 |
| *SPRR2E* | 0 | 0 | 1 | 0 |

Since YTHDF readers typically destabilize target transcripts, we next assessed the decay kinetics of representative RNAs. Following YTHDF1/2/3 silencing and 4 h *C. albicans* exposure, cells were treated with actinomycin D to halt new transcription, and target mRNAs were measured over a 4 h chase period. Half-life was determined by one-phase exponential decay modelling. Knockdown of YTHDF1/2/3 prolonged the stability of multiple transcripts (**Fig. 2b**), for example *CXCL8* and *CXCL10*. Thus, YTHDF proteins promote decay of transcripts, and hence their elevated steady-state abundance upon YTHDF knockdown reflects increased transcript stability rather than new transcriptional induction. YTHDF knockdown also increased secretion of CXCL8 protein, measured 24 h post-infection (**Fig. 2c**). Unexpectedly, *SPRR2A* and *SPRR2E* were also stabilized upon YTHDF depletion (**Fig. 2b**), despite their reduced overall expression (**Fig. 1g-i**). Thus, for some transcripts effects beyond YTHDF-driven regulation contribute to overall regulation of mRNA expression.

In addition to controlling mRNA stability, YTHDF proteins can augment translation initiation of methylated transcripts ^59–61^. To determine whether YTHDF proteins alter translational activity in OECs upon *C. albicans* exposure, we evaluated global protein synthesis by monitoring puromycin incorporation in infected TR146 cells (2 and 4 h) subjected to YTHDF1/2/3 silencing. As shown, there was no detectable alteration in overall levels of ongoing protein synthesis following YTHDF knockdown in response to *C. albicans* (**Extended Data Fig. 2**).

### *C. albicans*-induced target transcripts are methylated *in vivo* during oral infection

While the TR146 OEC line is widely used to investigate the oral mucosal response to *C. albicans*, this system cannot capture the complexities of infection *in vivo*. Therefore, we next sought to assess changes in the epitranscriptomic landscape of the oral mucosa during oral *C. albicans* infection. We employed a standard model of oropharyngeal candidiasis (OPC) in which mice were infected sublingually with *C. albicans* ^62^. Wildtype (WT) (immunocompetent) mice typically clear the fungus within 3 days, and peak expression of inflammatory cytokines and chemokines in the oral mucosa (tongue) occurs at day 2 ^3^. Critical genes that coordinate the response to OPC in this time frame include SPRRs, chemokines, antimicrobial proteins and the Th17/IL-17 signaling pathway ^3,55,56,63–66^. First, we quantified m⁶A-modified RNA in tongues from uninfected mice (PBS/sham) compared to *C. albicans*-infected mice at 2 days post-infection ^3,63^. Total m⁶A levels in tongue were unchanged at this time point (**Fig. 3a**), indicating that *C. albicans* infection does not globally alter the epitranscriptome.

**Figure 3.**
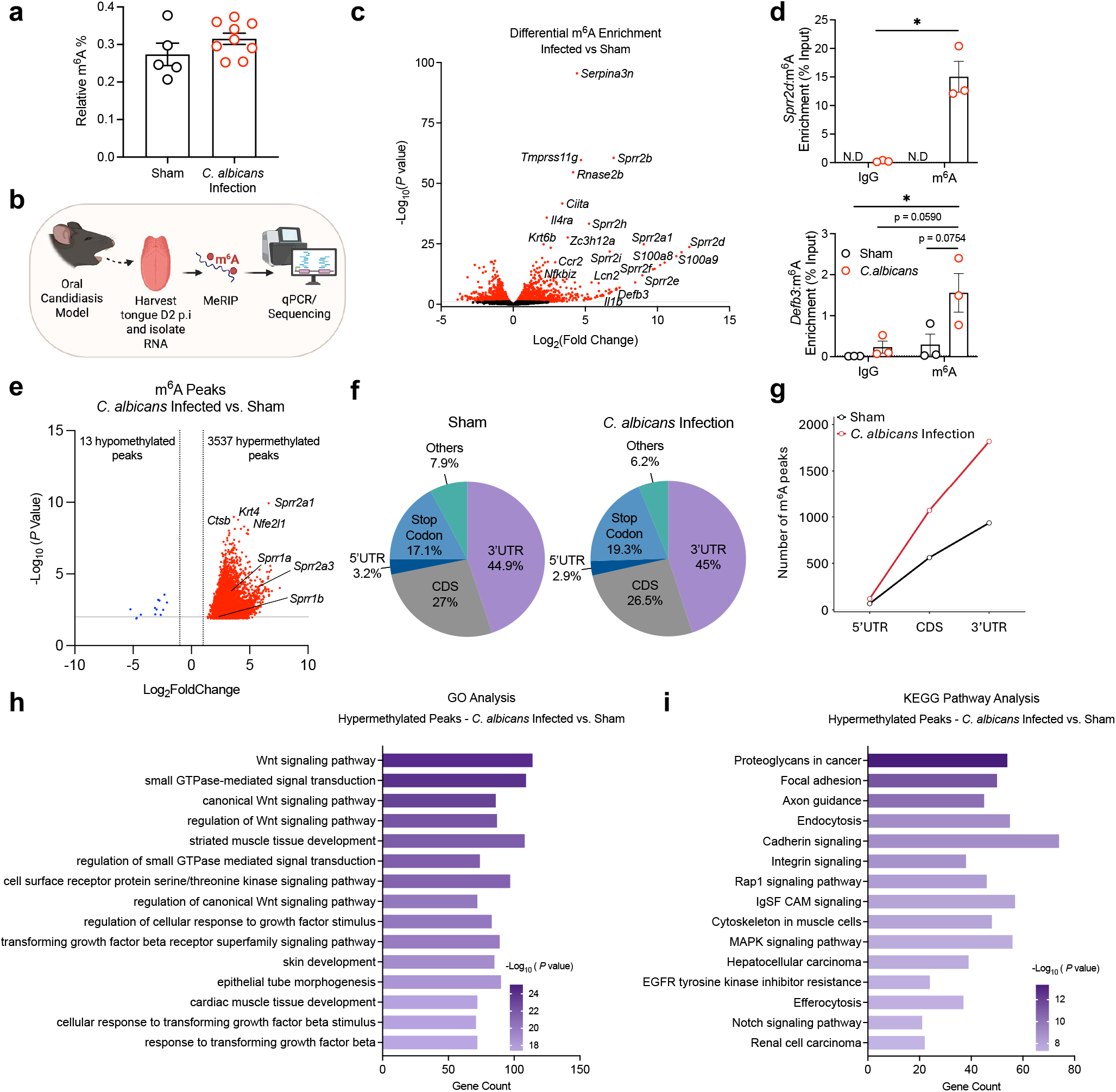
Oral *C. albicans* infection selectively reshapes the m^6^A landscape on infection-responsive genes. (**a**) Global m^6^A levels in sham (PBS) (black) and infected (red) murine tongue tissue at day 2 p.i. were measured by ELISA to detect m^6^A. Data show mean ± SEM (n=5–9 mice, 2 experiments). Significance was assessed by Welch’s t-test. (**b**) Workflow for m^6^A RNA immunoprecipitation (MeRIP). (**c**) WT mice were infected with *C. albicans,* and RNA from tongue was subjected to MeRIP. Volcano plot showing differentially enriched m^6^A-containing transcripts in the m^6^A RIP fraction from tongues of *C. albicans*-infected and sham-treated mice. Each point represents a transcript, plotted according to log_2_ fold change (infected vs. sham) and −log_10_ (*P* value). Significantly enriched transcripts are highlighted according to the indicated thresholds (adjusted *P* < 0.2 and |log_2_fold change| > 1). (**d**) Tongue RNA collected at 2 days p.i. was subjected to MeRIP (n= 3 mice per group). *Sprr2d* and *Defb3* were quantified by qPCR normalized to *Gapdh.* Significance was assessed by 2-way ANOVA with Bonferroni’s multiple comparisons test. (**e**) Volcano plot showing significantly hypermethylated or hypomethylated m^6^A peaks in total RNA of *C. albicans*-infected vs. sham/uninfected mice tongues. Each point represents an individual m^6^A peak plotted by log_2_fold change in methylation and −log_10_ (*P* value). Hypermethylated and hypomethylated peaks meeting the indicated significance thresholds (adjusted *P* < 0.05 and |log_2_fold change| > 1) are highlighted. The m^6^A-containing transcripts with notably increased (red) and decreased (blue) peaks are highlighted. (**f**) Proportion of m^6^A peak distribution in the 5′UTR, CDS, and 3′UTR regions across total transcriptome. (**g**) Profile of m^6^A peak distribution across the 5′UTR, CDS, and 3′UTR in sham or infected mice tongue. (**h**) GO and (**i**) KEGG pathway analysis of the genes that contained hypermethylated m^6^A peaks in *C. albicans*-infected mice tongue vs. sham/uninfected. Significance was assessed by 1-way ANOVA with Bonferroni’s multiple comparisons test.

To determine whether there were transcript-specific epitranscriptomic changes in the oral mucosa in response to infection, we performed MeRIP-seq on total RNA from infected and sham-treated tongues at day 2 (**Fig. 3b**). Differential expression analysis of the m^6^A enriched (RIP) fraction showed a substantial number of differentially enriched methylated transcripts (843 upregulated, 829 downregulated). Strikingly, these included many genes documented to be functionally associated with oral antifungal defense, inflammatory signaling, and antimicrobial responses, for example multiple *Sprr* genes (**Fig. 3c**) ^55,56^. IL-17 receptor signaling is essential for immunity to candidiasis ^3,67^ and in the context of autoimmunity IL-17 has been shown to signal through the m^6^A pathway ^40^. In the MeRIP of *C. albicans* infected tongue, *Il17a* was induced as expected but was not detectably methylated. However, many genes induced by the IL-17 pathway were identified in this screen, including *Defb3, S100a8/9, Serpins, Lcn2,* and *Zc3h12a* (**Fig 3c, d**). Consistent with their enrichment in MeRIP, all these genes contained SRAMP-predicted m^6^A modification sites (**Table 2, Extended Data Fig. 3**).

**Table 2.**
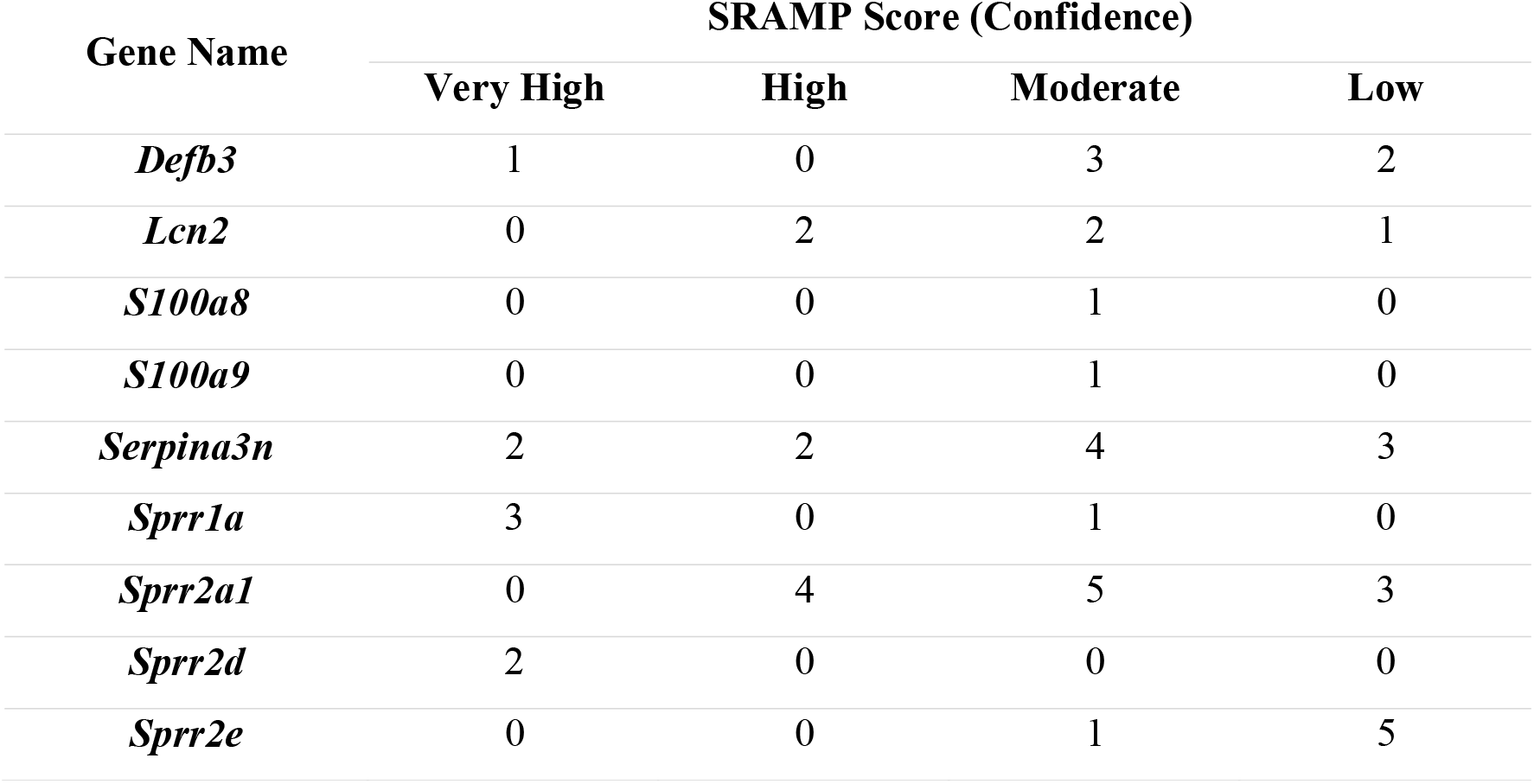
Predicted sites of m^6^A modification of murine AMPs identified using the SRAMP database.

Differential peak analyses of the m^6^A epitranscriptome across 6,121 detected peaks revealed a strong bias towards hypermethylation upon infection: 3,537 peaks were significantly hypermethylated following infection, compared to only 13 hypomethylated peaks (**Fig. 3e**). Top hypermethylated transcripts included *Sprr2a1*, as well as other genes not previously implicated in immunity to OPC, including *Ctsb*, *Krt4*, and *Nfe2l1* (**Fig 3e**, **Table 3)**, with multiple transcripts having more than one hypermethylated peak. To confirm that detected peaks represent authentic m⁶A modifications rather than technical artifacts, peaks were cross-referenced against input libraries, confirming enrichment on expressed transcripts and validating the high specificity of the m^6^A-RIP. This also confirmed that, even though total m^6^A levels did not change in the tongue, *C. albicans* infection drives widespread transcript-specific gains in m⁶A deposition. There was similarity in the total m^6^A distribution between sham and *C. albicans* infected tongues, which coincided with the typical m^6^A peak distribution features (**Fig. 3f**). Consistent with other studies ^25,68,69^, hypermethylated peaks were predominantly distributed across coding regions and 3′ UTRs (**Fig. 3f-g, Extended Data Fig. 4a**). Notably, infection with *C. albicans* resulted in elevated m^6^A peaks across the entire gene bodies compared to sham/uninfected (**Fig. 3g**).

**Table 3.**
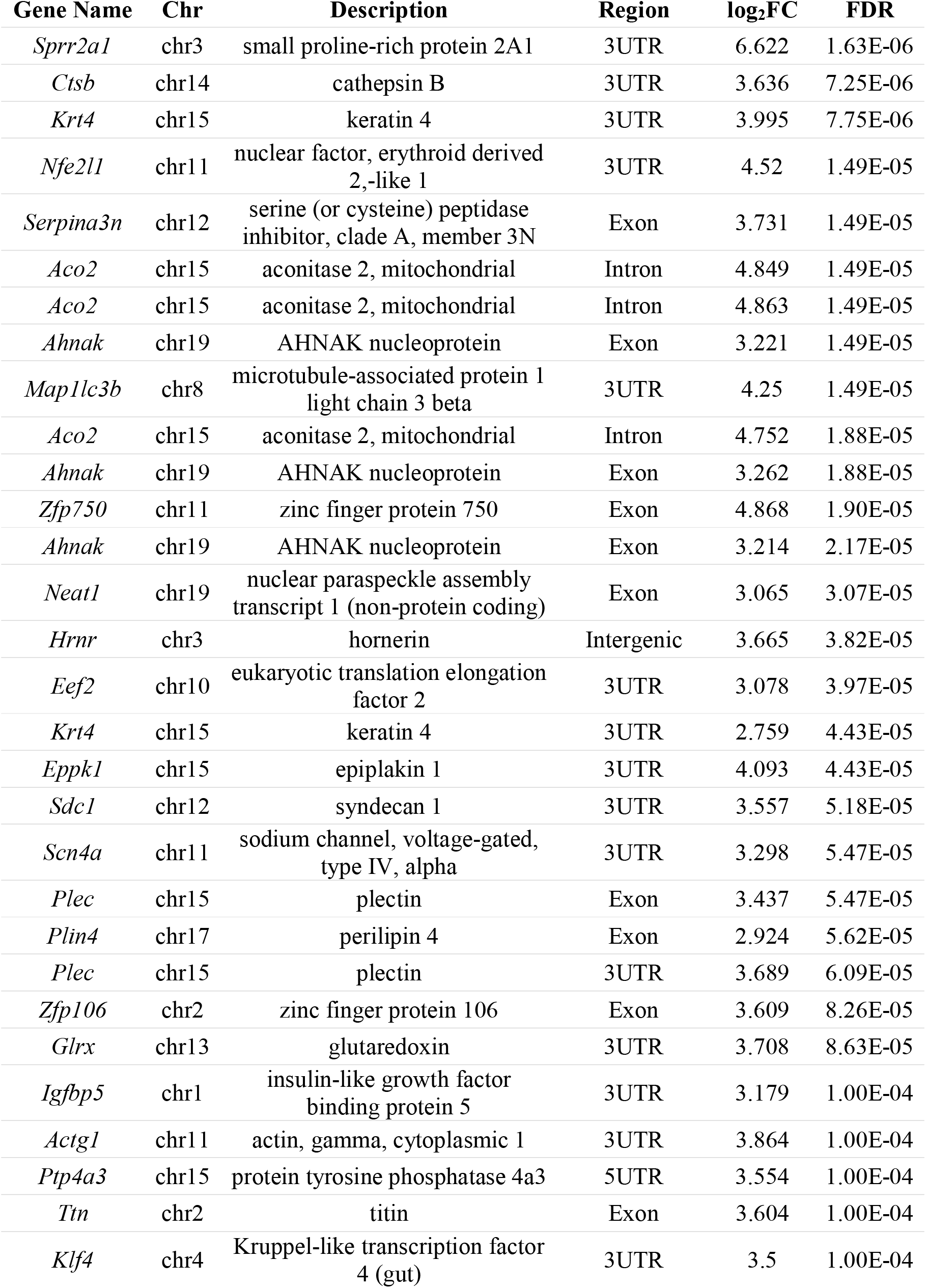
Top 30 differential m^6^A peaks in murine tongue tissue 2 days following *C. albicans* infection (ranked by FDR)

To identify the core infection-responsive m⁶A-regulated gene set, we examined the intersection of genes differentially expressed in the input fraction with those carrying hypermethylated peaks. This analysis revealed that 1,714 genes (50.5%) were hypermethylated without a corresponding transcriptional change in the input, while only 2 genes (*Sprr2a1*, *Tmprss119*) showed both transcriptional upregulation and gain of methylation (**Extended Data Fig. 4b-c**). A total of 444 genes were shared between the DEGs identified in the m⁶A RIP fraction and the hypermethylated peak set, representing transcripts that are both dynamically expressed and remodeled during infection. The majority of infection-associated methylation changes occurred on transcripts with stable abundance suggesting that RNA methylation is used primarily as a tuning mechanism to increase the effectiveness (stability or abundance) of already existing transcripts, leading to important post-transcriptional control of the epithelial response to oral *C. albicans* infection.

Enrichment analyses in KEGG pathways and GO ontology terms revealed that transcripts with increased m^6^A methylation peaks were preferentially over-represented in developmental signaling networks (Wnt, TGF-β and Notch pathways), as well as pathways regulating cell adhesion, cytoskeletal organization, epithelial morphogenesis, and small GTPase-mediated signaling (**Figure 3h-i**). Collectively, these pathways regulate epithelial architecture, tissue remodeling, and dynamic cellular signaling, suggesting that infection-induced m^6^A methylation preferentially targets transcripts involved in coordinating epithelial barrier adaptation and tissue responses during *C. albicans* infection.

Together, these data demonstrate that oral infection with *C. albicans* induces extensive and selective remodeling of the m⁶A epitranscriptome. Major targets of the m^6^A pathway are infection-responsive transcripts involved in antimicrobial defense and IL-17R signaling.

### Pharmacological METTL3 blockade ameliorates OPC

Current antifungal therapies are limited and compromised by development of drug resistance, highlighting the need for alternative therapeutic strategies ^70,71^. A recent study indicates that use of a pan-methyltransferase inhibitor, sinefungin, selectively reduces m^6^A modification within *Candida albicans* hyphae, suggesting that loss of fungal m^6^A modifications may be a mechanism underlying the antifungal capacity of this drug ^41^. Given that many of the m⁶A-modified transcripts we saw *in vivo* upon infection are known to control fungal pathogenesis ^3–5,50,63,72^, we hypothesized that blockade of epitranscriptomic machinery could constitute an unexplored therapeutic vulnerability during infection. To test this, we took advantage of a first-in-class METTL3 inhibitor, STM2457 ^73^ which has been shown to impair tumor growth in leukemia models ^73^ but has never been examined in fungal immunity. Mice were administered STM2457 every 1-2 days starting 5 days prior to oral infection with *C. albicans* (**Fig 4a**). STM2457 treatment reduced overall global m^6^A levels in the tongue, confirming its efficacy *in vivo* (**Fig. 4b**). At day 1, STM2457 substantially reduced oral fungal burdens compared to vehicle (23.9-fold reduction) (**Fig. 4c**). This trend continued at day 2, though the magnitude of reduction was less pronounced (**Extended Data Fig. 5a**). As is typical with this OPC model, infected control mice (vehicle-treated) lost weight in the early phase of infection, but mice treated with STM2457 maintained a stable body weight throughout the entire observation period, providing further evidence for a protective effect of METTL3 inhibition (**Fig. 4d**).

**Figure 4.**
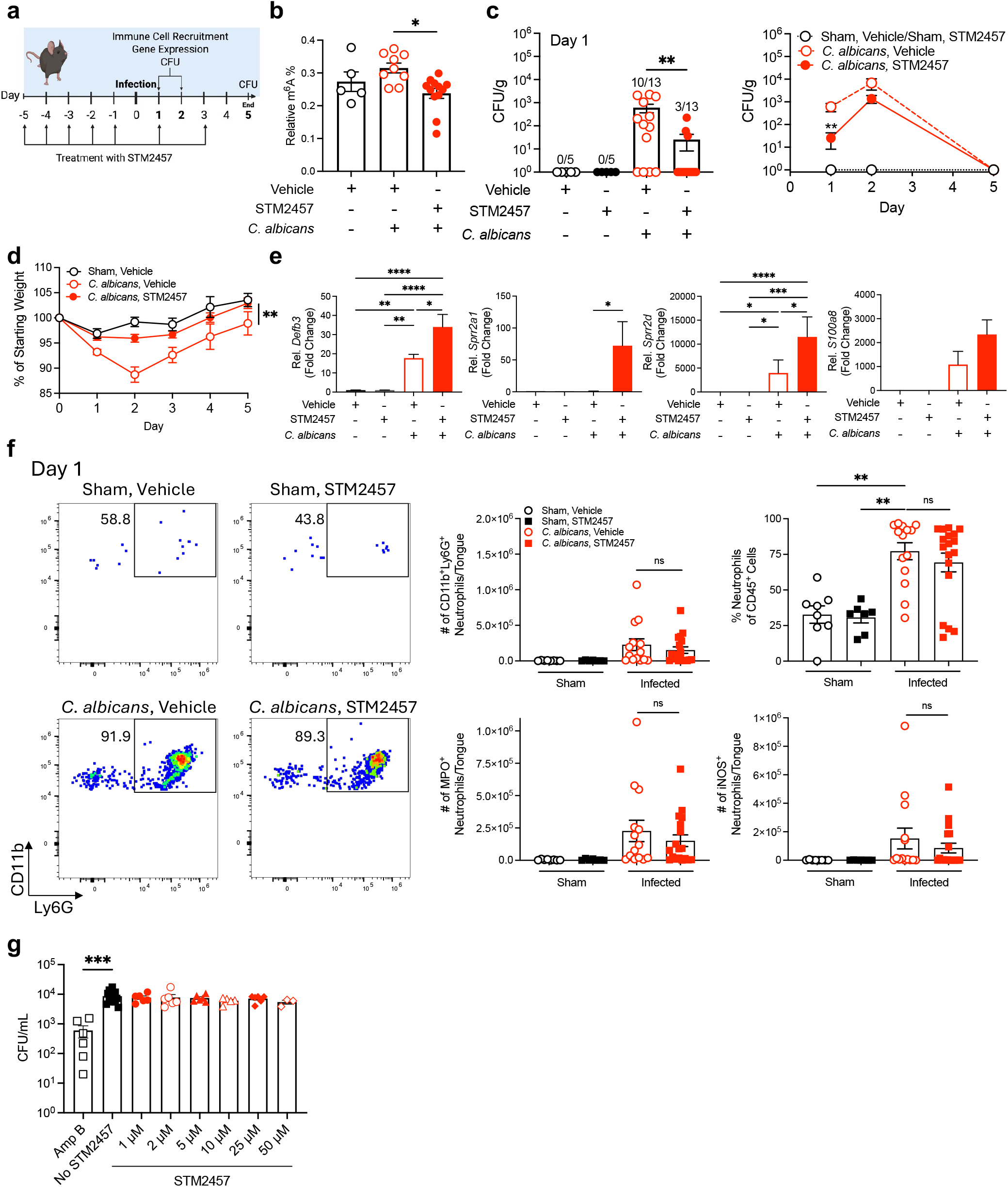
Inhibition of METTL3/m^6^A augments epithelial responses during oral *C. albicans* infection. (**a-c**) Indicated mice were treated with STM2457 via intraperitoneal injection on days −5 to −1 and infected with *C. albicans* or PBS (Sham). (**b**) Tissue m^6^A levels were measured in tongue on day 2; significance was assessed by Kruskal-Wallis with Dunn’s multiple comparisons test. (**c**) CFU in tongue on the indicated days post-infection. Data show mean of CFU/g tongue (n=3 independent experiments; 6–25 mice); significance was assessed by Mann-Whitney test. (**d**) Weight change over infectious time course ± STM2457 (n=6–25 mice); significance was assessed by 1-way ANOVA with Bonferroni’s multiple comparisons or Kruskal-Wallis with Dunn’s multiple comparisons test. (**e**) Gene expression in tongue on day 2, normalized to *Gapdh*; significance was assessed by 1-way ANOVA with Bonferroni’s multiple comparisons, Kruskal-Wallis with Dunn’s multiple comparisons test or Mann-Whitney test. (**f**) Neutrophil levels in tongue on day 1 p.i.. *Left*: Representative flow cytometry plot showing percent of CD11b^+^Ly6G^+^ neutrophils (gated on live CD45^+^ cells). *Right*: Pooled results (n= 4). Significance was assessed by Kruskal-Wallis with Dunn’s multiple comparisons test. (**g**) *C. albicans* yeast (SC5314) was cultured ± STM2457 for 2 h *in vitro* and CFU assessed by plating. Significance was assessed by Kruskal-Wallis with Dunn’s multiple comparisons test.

The oral host defense response to *C. albicans* is multifaceted, but antimicrobial peptides (AMPs) play essential roles, particularly β-defensin-3 ^5,67^ and β-defensin-1 ^74^. Other significant antifungal AMPs are SPRRs and calprotectin (S100A8/S100A9) ^55,56,72^. Indeed, blockade of METTL3 with STM2457 led to upregulation of β-defensin 3 (*Defb3*) and calprotectin (*S100a8)* during *C. albicans* infection ^10,75,76^ (**Fig. 4e**). Surprisingly, STM2457 treatment also upregulated SPRR genes (*Sprr2a1, Sprr2d*), which contrasted with findings in TR146 cells where knockdown of YTHDF1/2/3 decreased expression of the SPRR family (**Fig. 1**), indicating that SPRRs are potentially regulated by m^6^A readers beyond YTHDF1/2/3.

Myeloid cells are another important layer of protection against *Candida* and are recruited to the oral mucosa by OEC-derived chemokines ^3,64,67,76–78^ ^79^. However, the METTL3 inhibitor did not affect neutrophil recruitment or function (measured by MPO, iNOS) (**Fig. 4f, Extended Data Fig. 5b-c**). Similarly, METTL3 inhibition did not alter monocyte, macrophage, or T cell populations in the oral mucosa (**Extended Data Fig. 6).**

To determine whether the protective effects of STM2457 during OPC resulted from direct inhibition of *C. albicans* growth, we assessed fungal growth in the presence of increasing concentrations of STM2457. Treatment with STM2457 had no detectable effect on *C. albicans* growth across all concentrations tested (**Fig. 4g**), suggesting that the enhanced protection observed during OPC dominantly reflects modulation of host METTL3-dependent pathways rather than direct effects on the fungus. Together, these results suggest that the m^6^A pathway can be exploited as a therapeutic intervention in early responses to OPC, particularly through its influence on antifungal AMP gene regulation.

## Discussion

OECs comprise a key first line of defense against *C. albicans*, constituting a physical barrier and serving as active players in immune surveillance. Situated at a constantly renewing and microbially-exposed mucosal surface, OECs detect fungus, determine its capacity for ‘danger’ (i.e., hyphal formation), and orchestrate appropriate early innate immune responses ^80–85^. The signaling cascades that drive epithelial responses include PRR activation, NF-κB and AP-1 engagement, IL-17 and IL-22 cytokine signaling, all of which have been extensively characterized at the transcriptional level ^3–5,7,9,10,47,48,63,64,76,78,86–91^. Still, transcription represents only the first step in determining the magnitude of gene expression, as immune transcripts are highly prone to post-transcriptional regulation that allows for rapid and nuanced control of inflammatory responses. Here, we show that m^6^A modifications on *C. albicans*-induced mRNAs shape host antifungal responses. While the m^6^A pathway exerted both suppressive and enhancing effects on target genes, the net effect of blocking the m^6^A pathway pharmacologically was to improve early responses to *C. albicans*.

Particularly striking was the regulation of multiple AMPs such as β-defensins and SPRRs by m⁶A both *in vitro* and *in vivo*. While these are important effectors of oral antifungal defense ^3,55^, to our knowledge AMPs have not been previously shown to be regulated through epitranscriptomic mechanisms. SPRRs, originally characterized as markers of epidermal terminal differentiation, were recently shown to be antimicrobial effectors ^92^. Multiple SPRRs are induced during oral infection with *C. albicans* in an IL-17- and IL-22-dependent manner, and *Sprr1-3a*-deficient mice have an increased susceptibility to oral candidiasis. Sprr2a1 has direct candidacidal activity *in vitro* ^55^ and SPRR2e mitigated oral fungal burdens when applied therapeutically ^56^. β-defensins control *C. albicans* through direct fungicidal activity and may additionally act as endogenous alarmins and immune cell chemoattractants ^7,93^. While m^6^A-mediated restriction of AMP mRNA expression may somewhat restrict direct antifungal activity, this may be a means to prevent overexuberant inflammation. Therapeutically, m^6^A-mediated regulation of AMP genes is a potential target for boosting mucosal antifungal defense in OPC, aligning with findings that infection-induced remodeling of the epithelial m^6^A landscape modulates immune effectors ^44,46^.

Epitranscriptomic regulation of antifungal AMPs is part of a broader framework in which the m⁶A pathway serves as a homeostatic switch at barrier surfaces ^94–98^. For example, m^6^A can modulate intestinal responses, as knockdown of the m^6^A writer METTL14 in T cells caused spontaneous colitis, attributed to altered microbiome composition ^99,100^. Similarly, deficiency of the m^6^A eraser FTO reshaped the microbiome to increase anti-inflammatory *Lactobacillus* species over inflammatory-associated bacteria ^101^. By constraining inflammatory genes, m^6^A marks may aid in preserving tissue homeostasis while still allowing for rapid and effective responses when pathogenic overgrowth occurs. Administration of a METTL3 inhibitor during oral *C. albicans* infection enhanced expression of AMPs yet unexpectedly had no discernible effect on immune cell recruitment to the infected mucosa ^3,55^. The absence of changes in immune cell recruitment following METTL3 inhibition, despite a reduced fungal burden, is consistent with this interpretation and points to an epithelial-intrinsic defense mechanism of enhanced oral defenses.

Another surprising observation was that impairing m⁶A deposition was not functionally interchangeable with disrupting m⁶A recognition, exemplified by divergence between pharmacological METTL3 inhibition *in vivo* and YTHDF depletion *in vitro*. Notable in this regard were the *SPRR*s, where METTL3 inhibition produced opposite effects on gene expression compared to YTHDF silencing. Also discrepant was the finding that YTHDF1/2/3 silencing reduced total *SPRR* levels while still enhancing transcript half-life. It is possible that control of *SPRRs* by YTHDFs could occur through alternative readers or other YTHDF-mediated mechanisms. These findings may also point to limitations of the TR146 cell line as a discovery tool to define oral antifungal responses. While much of what is known about oral epithelial defense has come from this system, there are examples where immune factors strongly induced in TR146 cells are not predictive of their activities *in vivo* (for example, neither IL-6 nor CCL20 are critical to prevent OPC in mice ^102,103^. Here, we saw that many chemokines were strongly methylated and subject to control by YTHDFs, yet in murine OPC no chemokines were detectably m^6^A modified nor were there apparent defects in hematopoietic cell recruitment following METTL3 blockade.

The impact of m^6^A-RNA modifications during infections has been most widely studied in the context of viruses, where this mark modulates replication through modification of viral RNAs and regulation of host antiviral factors ^104^. In bacterial settings, METTL3/14-mediated m^6^A modification of TLR2/4 enhances receptor signaling ^18^, and YTHDF1-dependent translation of *TRAF6* regulates responses to *E. coli* ^42^. In intestinal epithelia, *Cryptosporidium* suppresses expression of the m^6^A eraser ALKBH5, reducing expression of key host defense genes ^46^. Our findings extend this to *C. albicans* infection, suggesting that m⁶A fine-tunes epithelial defenses. This is very relevant at the oral mucosa, where effective antimicrobial responses must be balanced against the need to tolerate *C. albicans* commensalism without excessive inflammation ^66,105^. Collectively, these observations emphasize that the effects of RNA methylation are context-dependent, with potential to either suppress or promote antimicrobial responses depending on the pathogen and immune pathways engaged.

Current antifungal drug options are hampered by toxicity and drug resistance ^70,71,106,107^, making the identification of new host-directed targets a priority. Inhibition of METTL3 with STM2457 in mice reduced fungal burden and prevented weight loss. Intriguingly, this improvement occurred without altering immune cell recruitment to the oral mucosa, suggesting that manipulation of epitranscriptomic pathways may preferentially augment direct antifungal defense proteins rather than broadly amplifying inflammation. STM2457 was developed as a first-in-class METTL3 inhibitor with efficacy in AML, impairing leukemic stem cell engraftment and prolonging survival in xenograft models ^108^. Our results identify RNA methylation machinery as an untapped therapeutic avenue for fungal infections. Notably, *C. albicans* redistributed m^6^A marks were located within a defined subset of infection-responsive transcripts rather than affecting methylation universally. This selectivity may also make pharmacological targeting of this pathway more feasible, without widespread or indiscriminate side effects.

Collectively, these findings establish epitranscriptomic regulation as a previously unappreciated determinant of defense against *C. albicans*. Rather than acting as a global regulator of gene expression, the m^6^A pathway in this context appears to somewhat selectively control the fate of fungal-responsive transcripts, enabling the oral mucosa to mediate antimicrobial defenses while avoiding excessive inflammatory responses. The observation that blockade of this pathway ameliorates preclinical signs of mucosal candidiasis provides a framework for exploring RNA methylation as a path for host-directed antifungal therapies.

## Methods

### Mice

Wild type mice (C57BL/6, both sexes aged 6-12 weeks) were from Jackson Laboratory or bred in-house. Mice were maintained in specific pathogen free conditions with food and water ad libitum and a 12-hour dark-light cycle. STM2457 in vehicle (5% DMSO, 40% PEG300, 5% Tween 80, 50% ddH2O (Selleckchem) or 20% (w/v) 2hydroxyprolpy β-cyclodextrin (GLPBIO)) was administered i.p. at 50 mg/kg 5 days before challenge. Mice were infected with *C albicans* by sublingual inoculation with cotton balls saturated with 10^7^ CFU *C. albicans* yeast (strain CAF2-1, cultured to log phase at 30°C) or PBS for 75 mins under general anesthesia ^62,67^. Weight was tracked daily and mice were euthanized if weight loss exceeded 25%. Tongue homogenates were prepared using a gentleMACS dissociator (Miltenyi Biotec) with C-tubes. Fungal burdens were determined by serial dilutions on YPD agar with ampicillin and cultured for 48 h at 30°C. Tongues were homogenized in Trizol using gentleMACS (Miltenyi Biotec) with M-tubes and RNA was isolated with RNeasy Mini Kits (Qiagen).

### Ethics Statement

All animal experiments were conducted according to national and international guidelines and approved by the University of Pittsburgh IACUC.

### Flow Cytometry

Tongue was minced and digested with collagenase IV (0.7 mg/mL) and 1x trypsin in HBSS for 30 mins. Cell suspensions were blocked with Fc receptor Abs (anti-CD16/CD32, Biolegend) and stained with the following antibodies for 20 mins at 4 °C (all antibodies were from Biolegend, unless noted) CD45.2-BV786, CD4-BUV395, CD8α-PE-Cy5, TCRβ-BUV496, TCRγδ-eF450, CD44-BV786, CD11b-AF700, CD11c-APC, Ly6C-BV570, Ly6G-PE-CF594, F4/80-PE. Cells were fixed and permeabilized with Cytofix/Cytoperm (BD Biosciences) and stained intracellularly for RORγt-BV650, Tbet-BV711, iNOS-PE-Cy5.5, or MPO-AF647 for 30 mins at 4 °C. Dead cells were excluded using the eF506 Viability Dye (eBiosciences). Flow cytometry was performed using a Cytek Aurora cytometer and analyzed using FlowJo software (FlowJo). Quantification was performed in FlowJo 10.9.0 and GraphPad Prism9.

### RNA Methylation and MeRIP (m^6^A-RIP) qPCR

Levels of m^6^A were quantified with an ELISA-based m^6^A RNA Methylation Quantification Kit (EpiGentek, #P-9005). MeRIP qPCR was performed according to a low-input m^6^A-seq protocol with modifications ^109^: 10-15 μg total RNA was fragmented (RNA Fragmentation Reagents, Thermo Fisher Scientific, AM8740) at 70°C for 15 min and purified by ethanol precipitation. 0.1 fmol of a control m^6^A-modified *Gaussia* luciferase RNA or unmodified *Cypridina* luciferase RNA (EpiMark *N6*-methyladenosine Enrichment kit) were spiked in all samples for normalization, with 10% of sample retained as input. For MeRIP, 25 μL protein A/G magnetic beads (Thermo Fisher Scientific) were incubated with anti-m^6^A antibody in 250 µL standard reaction buffer (10 mM pH 7.5 Tris-HCl, 150 mM NaCl, and 0.1% IGEPAL CA-630). Beads were washed twice with standard reaction buffer and incubated with the bead-antibody mixture at 4°C for 2 h. Samples were washed twice with reaction buffer, twice with low-salt reaction buffer (10 mM pH 7.5 Tris-HCl, 50 mM NaCl, and 0.1% IGEPAL CA-630), and twice with high-salt reaction buffer (10 mM pH 7.5 Tris-HCl, 500 mM NaCl, and 0.1% IGEPAL CA-630) for 10 min at 4°C. Bound RNA was eluted by competition with 6.7 mM N6-methyladenosine (Sigma) in 200 μL standard reaction buffer and purified with RNeasy MiniElute spin column, and suspended in RNase-free H_2_O. cDNA was synthesized with iScript cDNA synthesis Kit, and qPCR performed with SYBR Green Supermix. Transcripts were measured by qPCR in Input and m^6^A-IP RNA fractions. Ct values for RIP fractions were normalized to the corresponding Input fraction to account for RNA sample preparation differences: ΔCt [normalized RIP] = Ct [RIP] − (Ct [Input] − Log2[Input Dilution Factor]), where the Input Dilution Factor reflects the fraction of Input RNA saved (e.g., 6.644 cycles for a 1% Input fraction). Normalized RIP Ct values were averaged across replicates, and enrichment expressed as % Input = 2^(−ΔCt [normalized RIP]).

Putative m^6^A sites were determined using SRAMP (http://www.cuilab.cn/sramp) ^57,58^ that identifies m^6^A sites according to the DRACH consensus motif and flanking sequence features. Full-length mature sequences for the relevant mRNAs were obtained from NCBI and submitted in FASTA format using the mature mRNA model, and predicted sites across confidence categories (very high, high, moderate, and low confidence) were determined.

### Cell culture and RNA silencing

TR146 human buccal epithelial cells ^9,10,48,110,111^ were cultured in Dulbecco’s modified Eagle’s medium F-12 HAM (DMEM/F-12) (Gibco) supplemented with 15% FBS (Gibco) and 1% (v/v) penicillin–streptomycin) (Sigma–Aldrich). Serum-free DMEM/F-12 was used to replace normal growth medium 24 h before and during cell stimulations. TR146 cells were seeded in antibiotic-free media and transfected 18-24 h later with 50 nM siRNA (SMARTpool ON-TARGET plus, Dharmacon) in DharmaFect Reagent (Dharmacon). Culture media was replaced after 24 h, and stimulations occurred 24 h later. TR146 cells were infected with an MOI of 10 *C. albicans* yeast (SC5314).

### qPCR and RNA-Sequencing

RNA was isolated with RNeasy Mini Kits (Qiagen), cDNA was synthesized with iScript cDNA synthesis Kit, and qPCR performed with SYBR Green Supermix on a CFX Opus 96 (Bio-Rad). Data were normalized to *Gapdh*. Primers were from QuantiTect (Qiagen). For RNAseq, cDNA libraries were prepared from TR146 OECs stimulated with *C. albicans* (strain SC5314) for 4 h and RNASeq was performed on the Illumina NextSeq 500 platform by the Health Sciences Sequencing Core at the University of Pittsburgh. Sequencing reads were annotated and aligned to human hg19 using STAR. STAR alignment files were used to generate read counts for each gene and differentially expressed genes were determined using DeSeq2. RNASeq data was analyzed using Partek Flow Software. Upon acceptance, data will be made publicly available on GEO.

### MeRIP-Sequencing

Tongues were collected on day 2 p.i., and 15 μg of total RNA was used for (MeRIP) as described above. cDNA libraries were prepared, and RNA sequencing was performed on the Illumina NextSeq 500 platform by the Health Sciences Sequencing Core at the University of Pittsburgh. Raw sequencing reads were quality-checked with FastQC (v0.11.9), trimmed with Trimmomatic (v0.39), and aligned to the mouse genome (GRCm39/mm39, Ensembl release 112) using STAR (v2.7.11b; --outSAMtype BAM SortedByCoordinate, --outSAMattributes NH HI AS NM MD, --quantMode TranscriptomeSAM GeneCounts, --alignSJoverhangMin 8). Genome indices were constructed using the GRCm39 primary assembly FASTA with the Ensembl GRCm39.112 GTF annotation. Post-alignment quality was assessed with MultiQC (v1.30) and PCR duplicates were marked with Picard MarkDuplicates. Gene-level expression in Input samples was quantified using RSEM (v1.3.3; --paired-end, --estimate-rspd).

Differential gene expression between *C. albicans*-infected and uninfected mouse tongue was assessed using DESeq2 (v1.42), using RSEM counts imported via tximport (v1.30). Genes with fewer than 10 counts in at least the smallest group size were filtered prior to analysis. Significance was determined by Wald test with Benjamini-Hochberg correction; given limited sample size (n = 2–3/group), genes with adjusted p < 0.2 were considered differentially expressed. Ensembl IDs were converted to HGNC symbols using org.Mm.eg.db annotation package (v3.19).

Differential m^6^A methylation between infected and uninfected samples was identified using exomePeak2 (v1.16.2) and assessed by comparing m^6^A immunoprecipitation (IP) samples against matched Input samples using a negative binomial generalized linear model. A transcript database was constructed from the Ensembl GRCm39.112 GTF using *makeTxDbFromGFF* from the txdbmaker package and restricted to standard chromosomes (chromosomes 1–19, X, Y, MT). GC content bias correction was applied using the BSgenome.Mmusculus.UCSC.mm39 reference genome. The following parameters were used: strandness = “1st_strand” (confirmed by strand inference), test_method = “DESeq2”, fragment_length = 150 bp, bin_size = 25 bp, p_cutoff = 1×10⁻⁵, diff_p_cutoff = 0.05. Differential m^6^A peaks were annotated to the nearest gene using the *nearest* function from GenomicRanges (v1.54) with the TxDb.Mmusculus.UCSC.mm39.refGene annotation. Peaks were assigned to transcript regions - 5′UTR, CDS, 3′UTR, intron, or intergenic - based on overlap with features extracted from TxDb using GenomicFeatures (v1.54). Region assignments were applied in order of specificity: intron, exon, 5′UTR, 3′UTR, with 3′UTR taking precedence over the broader exon annotation. To validate that called peaks represent genuine m^6^A modification sites, sequences underlying hypermethylated peaks were extracted using *getSeq* from the BSgenome.Mmusculus.UCSC.mm39 package, centered on a 100 bp window around each peak summit. All 18 degenerate DRACH sequence variants (D = G/A/T, R = G/A, A, C, H = A/T/C) were searched using *vcountPattern* from Biostrings (v2.70). Sequence logos were generated using ggseqlogo (v0.2) from DRACH motif contexts extracted with 2 bp flanking sequence on each side.

Gene Ontology Biological Process (GO BP) and KEGG pathway enrichment analyses were performed using clusterProfiler (v4.10) with Benjamini-Hochberg correction and a significance threshold of adjusted p-value < 0.05. Reactome pathway analysis was performed using ReactomePA (v1.46). All pathway analyses used Entrez gene identifiers converted from gene symbols using the org.Mm.eg.db annotation database.

All analyses were performed in R (v4.6.0) with Bioconductor (v3.23). Figures were generated using ggplot2 (v3.5), EnhancedVolcano (v1.20), and ggseqlogo (v0.2). The full pipeline is available on Github and available upon request.

### RNA Decay Analysis

Following *C. albicans* infection, cells were washed with DMEM/F12 to remove non-adherent fungi and transcription was inhibited by actinomycin D (5 μg/mL; Sigma-Aldrich). Total RNA was isolated at the indicated times using RNeasy Mini Kits (Qiagen). cDNA was synthesized using the iScript Reverse Transcription Supermix for RT-qPCR (Bio-Rad), and transcript abundance quantified by RT-qPCR. Relative mRNA levels were normalized to the reference gene and expressed as the percentage of mRNA remaining relative to the time of actinomycin D addition (t = 0). RNA decay kinetics were analyzed by nonlinear regression using a one-phase exponential decay model in GraphPad Prism (v10), and transcript half-lives were calculated from the resulting decay curves.

### Immunoblotting, ELISA

TR146 cells were seeded in DMEM-F12 24 hours before transfection. Following transfection for 48 h, cells were stimulated with *C. albicans* for 2 or 4 h, washed and pulsed with 1 μg/mL puromycin (Sigma-Aldrich) at 37°C. Total cell lysates were subjected to SDS-PAGE and immunoblotted with anti-puromycin (Millipore) or anti-β-actin (Abcam).

Conditioned cell culture supernatants were collected 24 h post stimulation and ELISAs (R&D Systems) were performed following the manufacturer’s instructions.

### C. albicans Growth Assay

*Candida albicans* SC5314 was cultured in RPMI in the presence of vehicle (DMSO) or increasing concentrations of STM2457. At 2 h, serial dilutions were plated on YPD agar and incubated at 30°C for 24-48 h before CFU assessment.

### Statistics

All statistical analyses were done with GraphPad Prism software v11. using the D’Agostino and Pearson test, Anderson–Darling test, Shapiro–Wilk test and Kolmogorov–Smirnov test. Samples passing at least one normality test were considered to have normal distribution. Data were analyzed by 1-way ANOVA with Bonferroni’s multiple comparisons test (parametric data) or Kruskal-Wallis tests with Dunn’s multiple comparisons testing (nonparametric data). 2-way ANOVA with Bonferroni’s multiple comparisons test was used when comparing more than two sets of variables. Unpaired *t*-test or Mann-Whitney test was used for parametric or non-parametric datasets, respectively, when two groups were compared. *P* values < 0.05 were considered significant. Experiments were performed independently at least twice. Throughout, \**P* < 0.05, ** < 0.01, *** <0.001, ****<0.0001.

## Acknowledgments

SLG was supported by the University of Pittsburgh and the NIH (R37-DE022550). TCT and MEC were supported by T32-AI089443, and MEC by F32-AI186291. KY was supported in part by T32-AI074490 and F30-AI186235. RB was supported by the ANR-JCJC Program (ANR-22-CE15-0031) by the French National Research Agency and from the OI HEALTHI, HEADS and LSH Graduate School at Université Paris-Saclay. JD was supported in part by DP2-AI164325. The funders had no role in study design, data collection and analysis, decision to publish, or preparation of the manuscript. Work performed in the University of Pittsburgh Health Sciences Sequencing Core Facility services and instruments were supported, in part, by the University of Pittsburgh Office of the Senior Vice Chancellor for Health Sciences. We thank S. Jaffrey for valuable insights.

## Abbreviations

ALKBH: AlkB homologue
AMP: antimicrobial peptide
DEG: differentially expressed gene
FTO: fat and obesity associated protein
IGF2BP/IMP: insulin like growth factor mRNA binding protein
m^6^A: *N6*-methyladenosine
METTL: methyltransferase-like
MeRIP: m^6^A-RNA immunoprecipitation
OEC: oral epithelial cell
OPC: oropharyngeal candidiasis
RBP: RNA binding protein
RIP: RNA immunoprecipitation
SPRR: small proline-rich protein
YTHDF: YT521-B homology domain-containing family protein.

## Author Contributions

Conceptualization – SLG, NOP

Funding Acquisition - SLG, JD

Investigation – NOP, MEC, IM, ID, KY, TCT, BMC, SPV, RB

Visualization – NOP, IM, JD

Supervision – SLG, JD

Writing - Original draft – NOP

Writing - review and editing – SLG, JD, RB

## Supplementary Information

Ponde *et al.* BIORXIV_2025_67435

**Supplementary Data Figure 1: YTHDF reader proteins mediate *C. albicans*-induced inflammatory responses in OECs.** TR146 oral epithelial cells were transfected with siRNAs targeting YTHDF1, YTHDF2 or YTHDF3 individually. Cells were stimulated with *C. albicans* yeast 48 h for 4 h. Gene expression was quantified by qPCR normalized to *GAPDH*. Data are fold-change relative to siCtrl +PBS ± SEM (n= 2–3).

**Supplementary Data Figure 2: YTHDF reader proteins do not detectably impact global protein translation.** (A) Global translation measured by puromycin incorporation in control and YTHDF-depleted cells following infection. Puromycin-labeled nascent peptides were detected by immunoblotting with Abs against puromycine (top) or actin (bottom). (n=2)

**Supplementary Data Figure 3: Predicted m^6^A sites in *C. albicans* induced transcripts.** Predicted m^6^A sites in (**A**) *Sprr2d* and (**B**) *Defb3* were identified on the SRAMP platform. DRACH motifs are highlighted in red, splice regions in green and coding sequences in yellow.

**Supplementary Data Figure 4: Oral *C. albicans* infection in mice remodels m^6^A marks on infection-responsive genes.** WT mice (n=3) were infected orally with *C. albicans* SC5314 and on day 2 total RNA from tongue was subjected to MeRIP. (**A**) Hypermethylated m^6^A peaks in total RNA of *C. albicans*-infected vs. sham/uninfected mice tongues and corresponding m^6^A peak distribution in the 5′UTR, CDS, and 3′UTR regions across total transcriptome. (**B**) Overlap between DEGs identified in the input RNA, differentially enriched transcripts in the m^6^A RIP fraction, and hypermethylated transcripts identified by exomePeak2 following *C. albicans* infection. Numbers indicate overlap between each dataset. **(C**) Overlap analysis of transcriptional and m^6^A methylation changes following *C. albicans* infection.

**Supplementary Data Figure 5. METTL3/m^6^A suppression has no evident impact on neutrophil recruitment or function.** Indicated mice (n=9-25) were injected with STM2457 i.p. on days −5 to −1 and infected sublingually with *C. albicans* or PBS (Sham) on day 0. (**A**) Fungal load in tongue was measured at day 2 p.i. (**B**) Gating strategy for flow cytometry. (**C**) *Left*: Representative flow cytometry plot showing percent of CD11b^+^Ly6G^+^ neutrophils (gated on live, CD45^+^ cells). *Right*: Data from 4 experiments. Significance was assessed by 1-way ANOVA with Bonferroni’s multiple comparisons test.

Supplementary Data Figure 6. METTL3/m^6^A suppression does not impact on myeloid or leukocyte recruitment following oral *C. albicans* infection. Mice were injected with STM2457 i.p on days −5 to −1 and infected with *C. albicans* or PBS (Sham). Tongue homogenates were analyzed on day 1 and day 2. Cells were gated on live, CD45^+^ cells and indicated immune populations gated as shown (N= 4 independent experiments). **(A**) Gating strategy of flow cytometry data acquired at day 1 p.i.. (**B**) Quantification of tongue immune cells treated ± STM2457. Significance was assessed by 1-way ANOVA with Bonferroni’s multiple comparisons test

