## Supplementary figures and images for "N6-methyladenosine (m^6^A) RNA modification restrains antifungal immunity and is a therapeutic target in oral candidiasis"

### fig s1

#### Extended Data Fig. 1

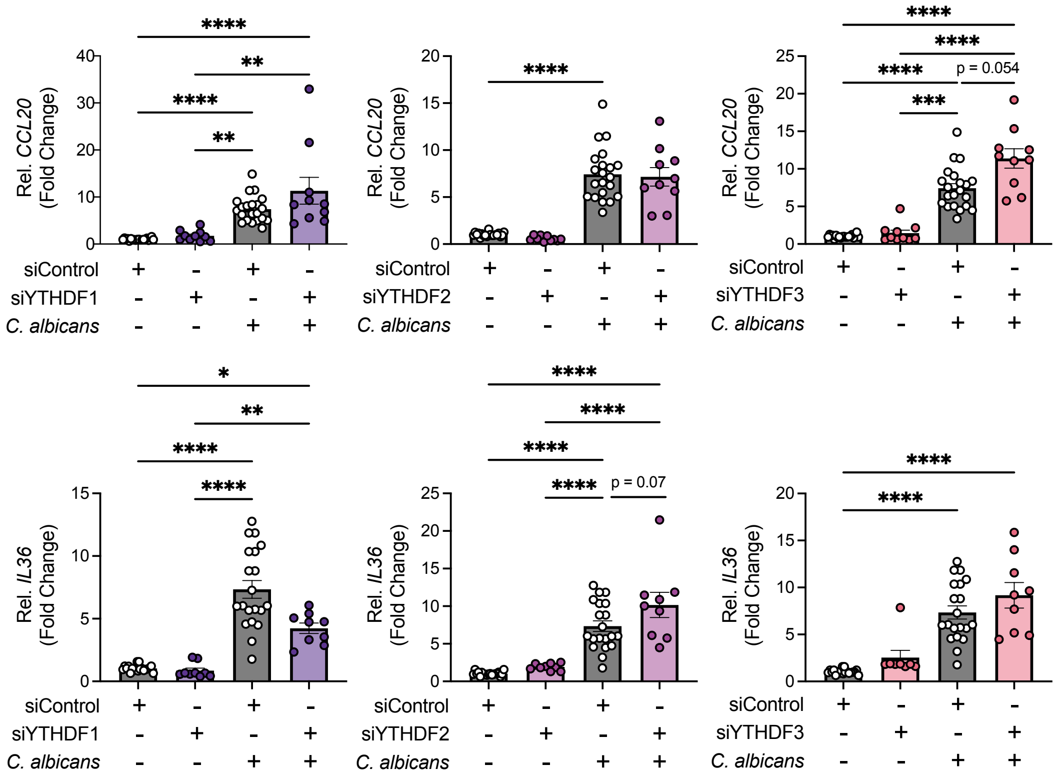

### fig s2

#### Extended Data Fig. 2

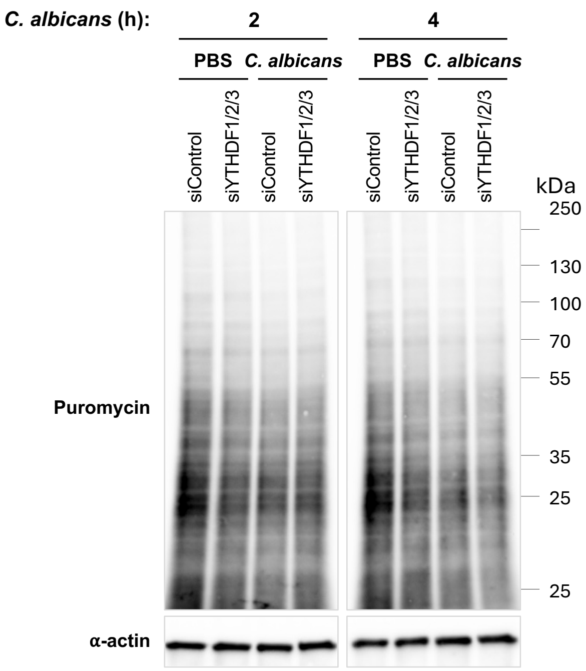

### fig s4

#### Extended Data Fig. 4

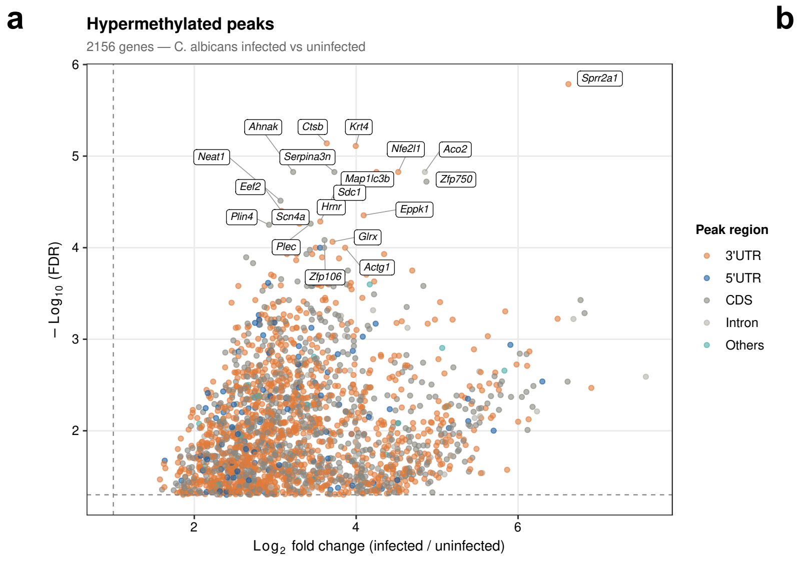


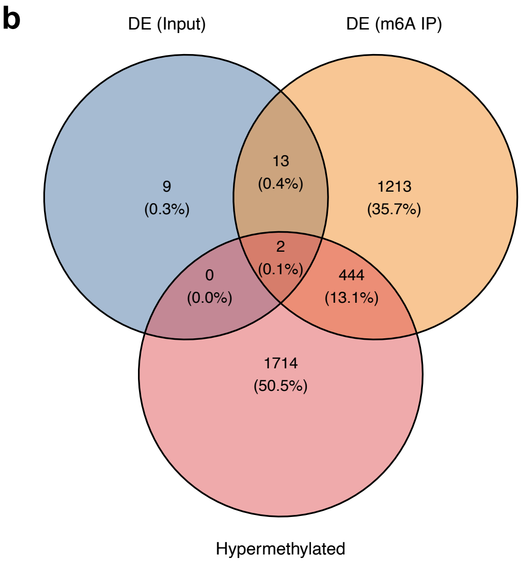


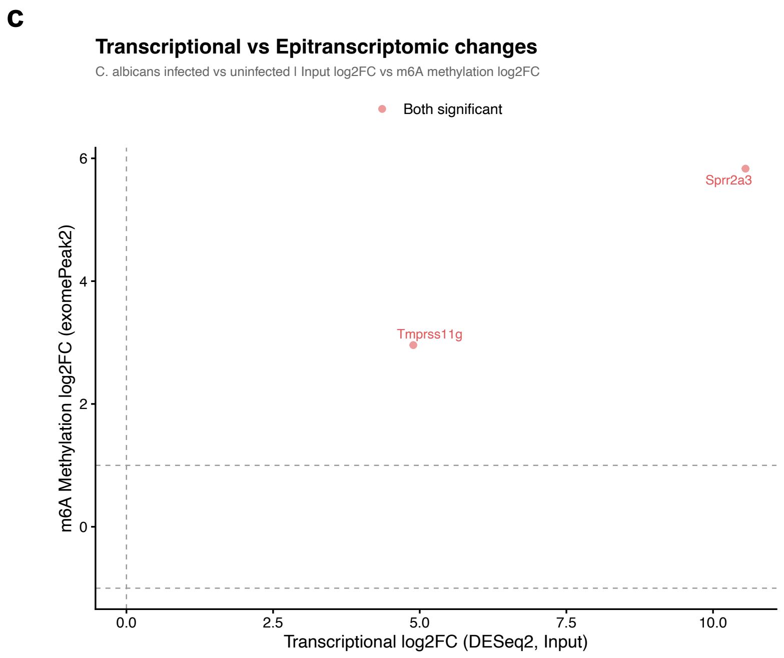

### fig s5

##### Extended Data Fig. 5

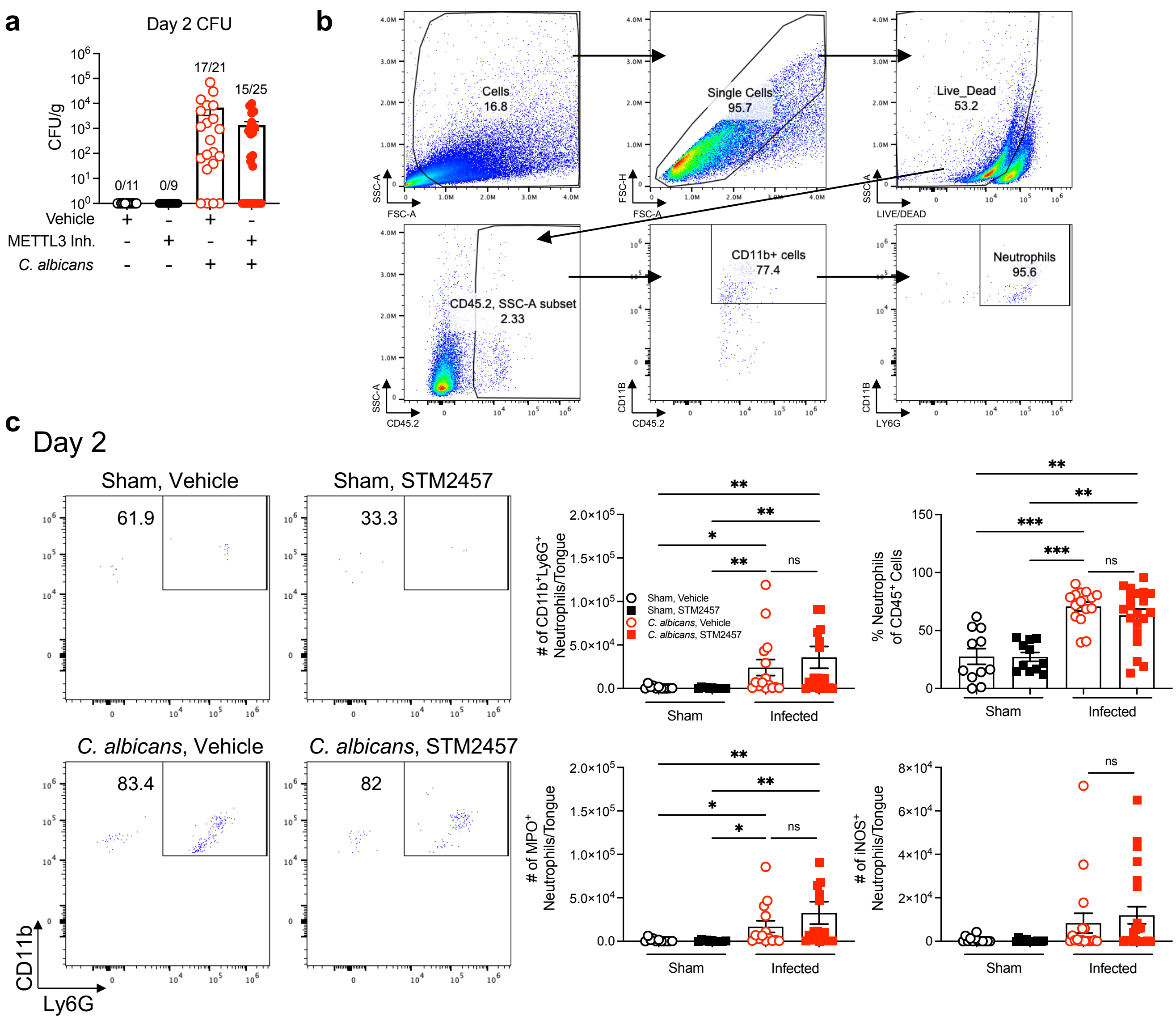

### fig s6

##### Extended Data Fig. 6

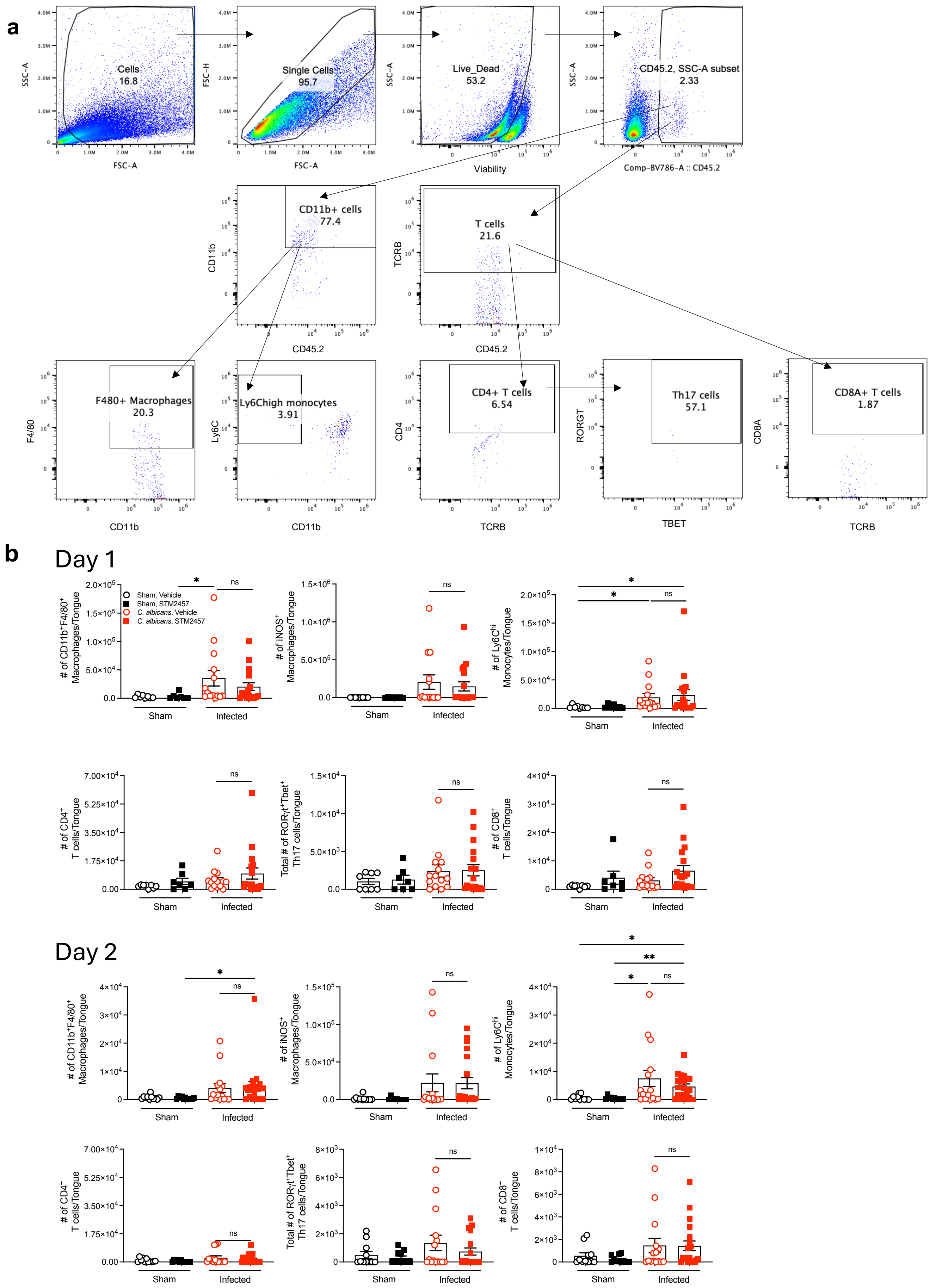
