## Supplementary material for "N6-methyladenosine (m^6^A) RNA modification restrains antifungal immunity and is a therapeutic target in oral candidiasis": fig s3

### Extended Data Fig. 3

## a

**NM_013756.2 Mus musculus defensin beta 3 (Defb3), mRNA to ENSMUST00000033852**

Splice CDS DRACH Known m6A

```
Ref.        1  CCAGGCTTCAGTCATGAGGATCCATTACCTTCTGTTTGCATTTCTCCTGGTGCTGCTGTC   60
Query          -CAGGCTTCAGTCATGAGGATCCATTACCTTCTGTTTGCATTTCTCCTGGTGCTGCTGTC
Prob.

Ref.       61  TCCACCTGCAGCTTTTAGCAAAAAAATCAACAATCCAGTAAGTTGTTTGAGGAAAGGAGG   120
Query          TCCACCTGCAGCTTTTAGCAAAAAAATCAACAATCCAGTAAGTTGTTTGAGGAAAGGAGG
Prob.

Ref.      121  CAGATGCTGGAATCGGTGCATTGGCAACACTCGTCAGATTGGCAGTTGTGGAGTTCCTTT   180
Query          CAGATGCTGGAATCGGTGCATTGGCAACACTCGTCAGATTGGCAGTTGTGGAGTTCCTTT
Prob.

Ref.      181  CCTCAAATGCTGCAAGAGAAAATAGAGAAGACCAAGATCCCGTTAAACAAAGAAAACACC   240
Query          CCTCAAATGCTGCAAGAGAAAATAGAGAAGACCAAGATCCCGTTAAACAAAGAAAACACC
Prob.

Ref.      241  AGAATTTGCTCCTCCATGAAGATGGACAAATCTTA-------------------------   300
Query          AGAATTTGCTCCTCCATGAAGATGGACAAATCTTAAACCAAATTAATATTCCTTGATTAA
Prob.                                  93%

Ref.      301  --------------------------------------------------           349
Query          ATGCAAACAAGTGCTCCCTAGCAAAAAAAAAAAAAAAAAAAAAAAAAAAA
Prob.
```

## b

**NM_011470.3 Mus musculus small proline-rich protein 2D (Sprr2d), mRNA to ENSMUST00000047477**

Splice CDS DRACH Known m6A

```
Ref.        1  AGACTCCTGGTACTCAAGGCCGAGACTACTTTGGAGAACCCGATCCTGAGAATCCAGCAC   60
Query          AGACTCCTGGTACTCAAGGCCGAGACTACTTTGGAGAACCCGATCCTGAGAATCCAGCAC
Prob.            0%                    0%           0%

Ref.       61  TATGTCTTACCAGCAGCAGCAGTGCAAGCAGCCTTGCCAGCCTCCACCTGTGTGCCCACC   120
Query          TATGTCTTACCAGCAGCAGCAGTGCAAGCAGCCTTGCCAGCCTCCACCTGTGTGCCCACC
Prob.

Ref.      121  CAAGAAGTGCCCTGAGCCTTGTCCTCCTCTAAAGTGTCCTGAGCCTTGTCCTCCACCAAA   180
Query          CAAGAAGTGCCCTGAGCCTTGTCCTCCTCTAAAGTGTCCTGAGCCTTGTCCTCCACCAAA
Prob.

Ref.      181  GTGCCCTGAGCCTTGTCCTCCTCCAAAATGCCCAGAGCCTTGTCCTGAGCCATGTCCCCC   240
Query          GTGCCCTGAGCCTTGTCCTCCTCCAAAATGCCCAGAGCCTTGTCCTGAGCCATGTCCCCC
Prob.

Ref.      241  TCCCTCATGCCAGCAGAAATGCCCTCCTGCGCAACCTCCTCCACCCTGCCAGCAGAAGTG   300
Query          TCCCTCATGCCAGCAGAAATGCCCTCCTGCGCAACCTCCTCCACCCTGCCAGCAGAAGTG
Prob.

Ref.      301  CCCACCTAAGAGCAAGTGAGGTCTTCAGCAGTCATCAGGACAAAGGGAAAAGAAGAGAAT   360
Query          CCCACCTAAGAGCAAGTGAGGTCTTCAGCAGTCATCAGGACAAAGGGAAAAGAAGAGAAT
Prob.                                                 0%

Ref.      361  CTATCTCATGTACTTCCAAAGCAACACCCATCTTCCTTCCAAATCCTGGCATGATTGCAG   420
Query          CTATCTCATGTACTTCCAAAGCAACACCCATCTTCCTTCCAAATCCTGGCATGATTGCAG
Prob.

Ref.      421  AGGAATCTCCTTCACCTTTCACCCTCCATGTTCCTGTGATGTCCCCTAACAGGGAAGATG   480
Query          AGGAATCTCCTTCACCTTTCACCCTCCATGTTCCTGTGATGTCCCCTAACAGGGAAGATG
Prob.                                                          0%

Ref.      481  TTTCCCCGAAGTTGTCCTGCTTTTGGTGTCAAGGGGCTGCAGAGAATGATCTGTCCTCAG   540
Query          TTTCCCCGAAGTTGTCCTGCTTTTGGTGTCAAGGGGCTGCAGAGAATGATCTGTCCTCAG
Prob.

Ref.      541  TTATTCAGTATCCCAGGTGCTCAGTGGGAACTATAGCTGCTATCTATCCTGTGCCCTCAG   600
Query          TTATTCAGTATCCCAGGTGCTCAGTGGGAACTATAGCTGCTATCTATCCTGTGCCCTCAG
Prob.                                      94%

Ref.      601  GAGATGTTGCCAAAAACTCTCAATCACCTTTGGTTTTGTGTTGTCAGCATGGTTTCATTT   660
Query          GAGATGTTGCCAAAAACTCTCAATCACCTTTGGTTTTGTGTTGTCAGCATGGTTTCATTT
Prob.                        93%

Ref.      661  CCTGAATAAAGTACAATCTAC-------                                  689
Query          CCTGAATAAAGTACAATCTACATAATGA
Prob.
```
